# Systemic mutagen exposures reported by normal kidney cell genomes

**DOI:** 10.64898/2026.04.07.716715

**Authors:** Yichen Wang, William Knight, Aida Ferreiro-Iglesias, Behnoush Abedi-Ardekani, My H. Pham, Sarah Moody, Yvette Hooks, Federico Abascal, Charles Nunn, Stephen Fitzgerald, Thomas Cattiaux, Valérie Gaborieau, Akihiko Fukagawa, Viorel Jinga, Stefan Rascu, Cristian Sima, David Georgievich Zaridze, Anush Felixovna Mukeria, Ivana Holcatova, Anna Hornakova, Naveen S. Vasudev, Rosamonde E. Banks, Simona Ognjanovic, Slavisa Savic, Maria Paula Curado, Stênio de Cássio Zequi, Rui Manuel Reis, Wesley Justino Magnabosco, Fernanda Vianna, Brasil Silva Neto, Sonata Jarmalaite, Algirdas Zalimas, Lenka Foretova, Marie Navratilova, Larry Phouthavongsy, Carolyn Shire, Worapat Attawettayanon, Surasak Sangkhathat, Chuling Ding, Andrew R. J. Lawson, Calli Latimer, Laura Humphreys, Peter J. Campbell, Sandra Perdomo, Iñigo Martincorena, Ludmil B. Alexandrov, Sam Behjati, Kourosh Saeb-Parsy, Tatsuhiro Shibata, Paul Brennan, Michael R. Stratton

## Abstract

Lifestyle, environmental and other exposures to exogenous mutagens generate somatic mutations in normal human cells *in vivo* and increase cancer risk. However, the global repertoire of exogenous mutagen exposures is uncertain. The mutational signatures of mutagens in normal tissues offer opportunities to detect such exposures and survey them at population level. Using single-molecule duplex sequencing of normal kidney (n=319) and blood (n=272) samples from 10 countries, we show that normal kidney cell genomes report an extensive repertoire of somatic mutational signatures. Microdissection of kidney structures revealed that proximal tubules exhibit higher mutation rates than other components of the nephron and most normal cell types despite low cell division rates. This is explained by marked enrichment of mutational signatures due to known exogenous carcinogenic mutagens including the plant-derived aristolochic acids, as well as several signatures of unknown causes including an unknown agent prevalent in Japan (SBS12), and signatures of uncertain origins (SBS40b and SBS40c). The results suggest the existence of multiple, common, systemically circulating mutagens affecting human populations and indicate that the genomes of kidney proximal tubule cells report such exposures with high sensitivity.

## Introduction

International cancer epidemiology has demonstrated geographical differences in the incidence rates of most common adult solid cancers^1^ which are likely predominantly due to differences in environmental and lifestyle exogenous exposures. Approximately 80% of cancer incidence in high-income countries is estimated to arise from such exposures, with about half attributable to known carcinogenic influences including tobacco smoking, obesity, and infectious agents^2–5^. Identifying the causes of the remainder could provide further opportunities for cancer prevention. However, few causes of cancer have been discovered in recent decades and approaches complementary to conventional epidemiology may be expedient in detecting them^6^.

Many known carcinogens are mutagens that cause distinct patterns of mutations, known as “mutational signatures”, in the genomes of exposed cells^7,8^. Cataloguing mutational signatures in human cells thus provides a record of lifetime mutagen exposures and constitutes a complementary approach to detecting some carcinogenic influences. Normal cells at body surfaces exposed to exogenous mutagens, for example the skin epidermis (ultraviolet light)^9^ and bronchial epithelium (tobacco smoke chemicals)^10^, exhibit high burdens of mutational signatures due to the agents they are directly exposed to and elevated rates of cancer consequent on these exposures. However, many exogenous mutagens are ingested, absorbed and circulated systemically by the blood, exposing multiple internal organs. Such mutagens, including aristolochic acids, aflatoxins and tobacco smoke chemicals, confer increased cancer risks but these are often limited to certain tissue or cell types. The basis of this specificity is not well understood.

The Mutographs Cancer Grand Challenge has recently sequenced the genomes of ∼5,000 cancers of eight organs from more than 30 countries to investigate geographical differences in cancer incidence and the exposures underlying them^11^. The study of clear cell kidney cancer genomes revealed multiple mutational signatures in addition to those of ubiquitous endogenous mutational processes, some of which showed geographical variation^12^. Interpretation of this observation, is, however, complicated because cancer genomes often acquire mutations caused by endogenous mutational processes absent in normal cells during neoplastic transformation^13^, rendering it uncertain whether some mutational signatures are endogenous or exogenous in origin. Exogenous mutagens are, however, expected to initially generate mutations in exposed normal cells. Hence, an important criterion favoring the exogenous origin of a signature is its presence in normal cells. In this study, therefore, we have applied recently developed single-molecule duplex DNA sequencing technology, which detects somatic mutations in multiclonal normal tissues, to normal kidney cell genomes to ascertain mutational signatures potentially generated by systemically circulating exogenous mutagens and to compare the mutation burdens conferred in different cell types.

## Results

### Single-molecule duplex sequencing of normal kidneys

Normal kidney samples from 319 individuals were used to investigate the mutational processes operative in normal human kidney cells (Fig.1a–b, Supplementary Table 1). 303 donors from 10 countries who had participated in Mutographs had clear cell renal cell carcinoma (ccRCC): Brazil (*n* = 40), Canada (*n* = 7), Czechia (*n* = 68), Japan (*n* = 66), Lithuania (*n* = 11), Romania (*n* = 33), Russia (*n* =37), Serbia (*n* = 13), Thailand (*n* = 4), UK (*n* = 24), with ccRCC whole genome sequencing data available for 296 of them. 16 additional UK donors had no history of renal cancer. DNAs extracted from normal kidney cortex (referred to as “bulk” samples to distinguish them from microdissected samples described below) were sequenced to detect somatic mutations using NanoSeq^14^, a highly error-corrected form of duplex DNA sequencing which enables reliable detection of somatic mutations in multiclonal cell populations. After library preparation, sequencing and decontamination, bulk sequencing data was obtained from 299 individuals (Supplementary Table 2). Peripheral blood samples from 272 Mutographs ccRCC patients (including 217 normal kidney donors) were also sequenced using NanoSeq (Supplementary Table 3) to compare somatic mutation burdens and mutational signatures in normal blood and normal kidney.

**Fig. 1.**
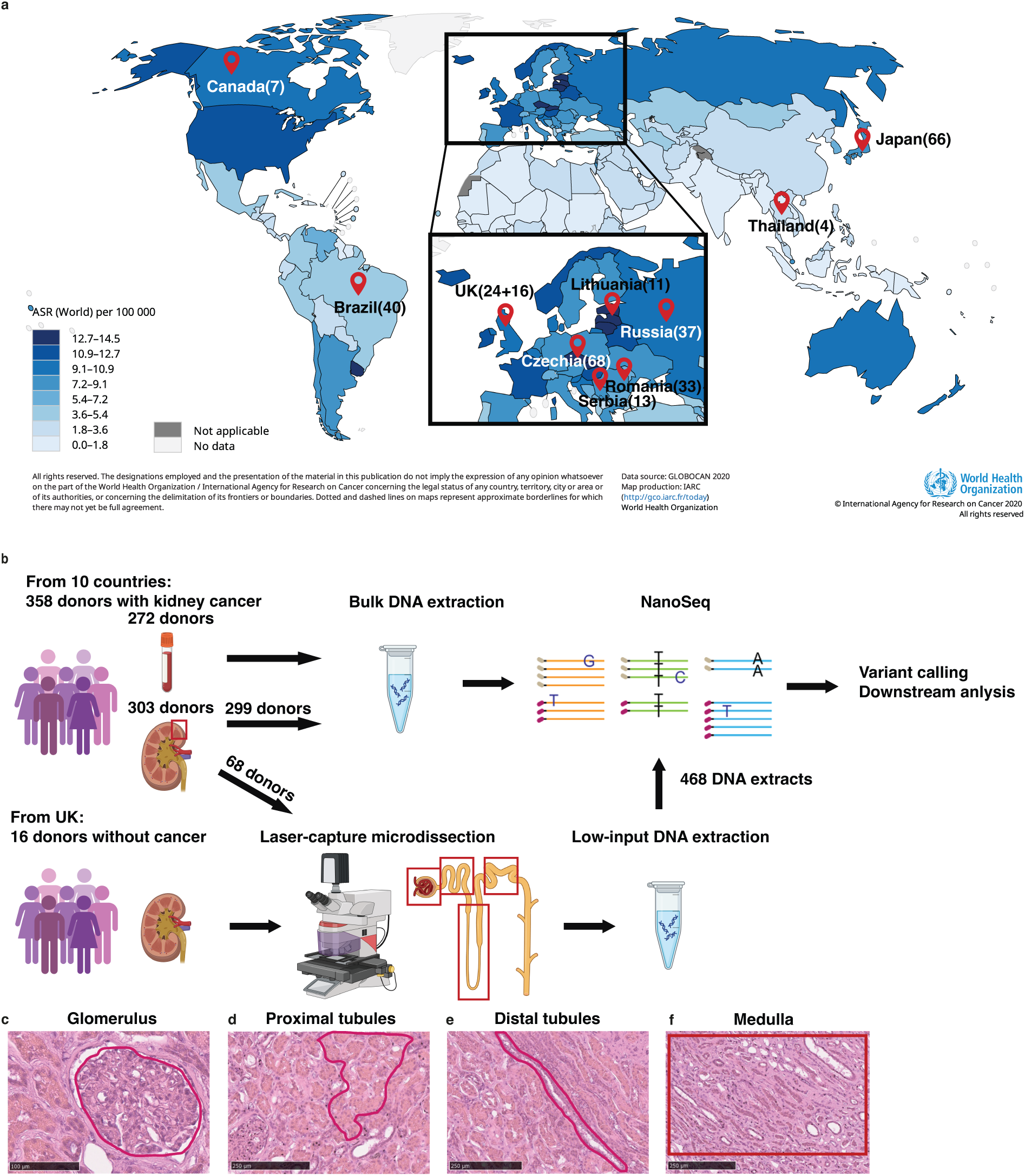
Study design. **a**, Geographic distribution of the 303 ccRCC patients and 16 non-cancer donors across 10 countries. ASR: age-standardized incidence rate of kidney cancer, for men and women combined. Data from GLOBOCAN 2020. Markers indicate countries included in this study (number of participants per country). **b,** Schematic workflow of the study. Modified from illustration created with BioRender.com. **c–f**, Illustration of H&E stained parts of the nephron for laser-capture microdissection.

The functional unit of the kidney, the nephron, is constituted of distinct, microscopically visible structural components which sequentially include the glomeruli, proximal tubules, medullary tubules and distal tubules. To analyze these separately for their mutation burdens and signatures, pooled sets of laser-capture microdissected (LCM) glomeruli, proximal tubules, medulla and distal tubules (Fig.1c) from each of 84 individuals (68 with ccRCC from the 10 countries and 16 with no renal cancer from UK) were sequenced for somatic mutations using NanoSeq (in total, 468 libraries; 173 from glomeruli, 166 from proximal tubules, 16 from distal tubules, and 113 from medulla) (Supplementary Table 4). Standard 20-fold coverage whole-genome sequencing data was generated from either blood or normal kidney DNA from all individuals to catalogue inherited variation and enable identification of somatic mutations. Samples were reviewed by a histopathologist with expertise in kidney disease diagnosis.

### Mutational signatures in normal kidney cells

The average total single-base substitution (SBS) and small insertion and deletion (indel) mutation burdens per country in diploid cell genomes from normal bulk kidney samples showed substantial geographical variation, similar to that previously reported in ccRCC^12^, with approximately two-fold higher burdens in Romania and Serbia compared to the other countries studied (Fig.2a–b and Extended Data Fig.1). To investigate the mutational processes underlying this variation, all sequenced bulk kidney cortex and laser-capture microdissected samples were included in *de novo* mutational signature extractions by two different algorithms (HDP based on a hierarchical Dirichlet process^15^ and SigProfilerExtractor based on non-negative matrix factorization^16^). The extracted SBS signatures (Extended Data Fig.2–3) were decomposed into COSMIC v3.4 reference signatures^17^ or, if not achievable, designated as new signatures (materials and methods). This resulted in the identification of COSMIC reference signatures SBS1, SBS2, SBS5, SBS12, SBS13, SBS22a, SBS22b, SBS40a, SBS40b, SBS40c and non-reference signatures SBSA, SBSB, SBSC and SBSD (Fig.2c).

**Fig. 2.**
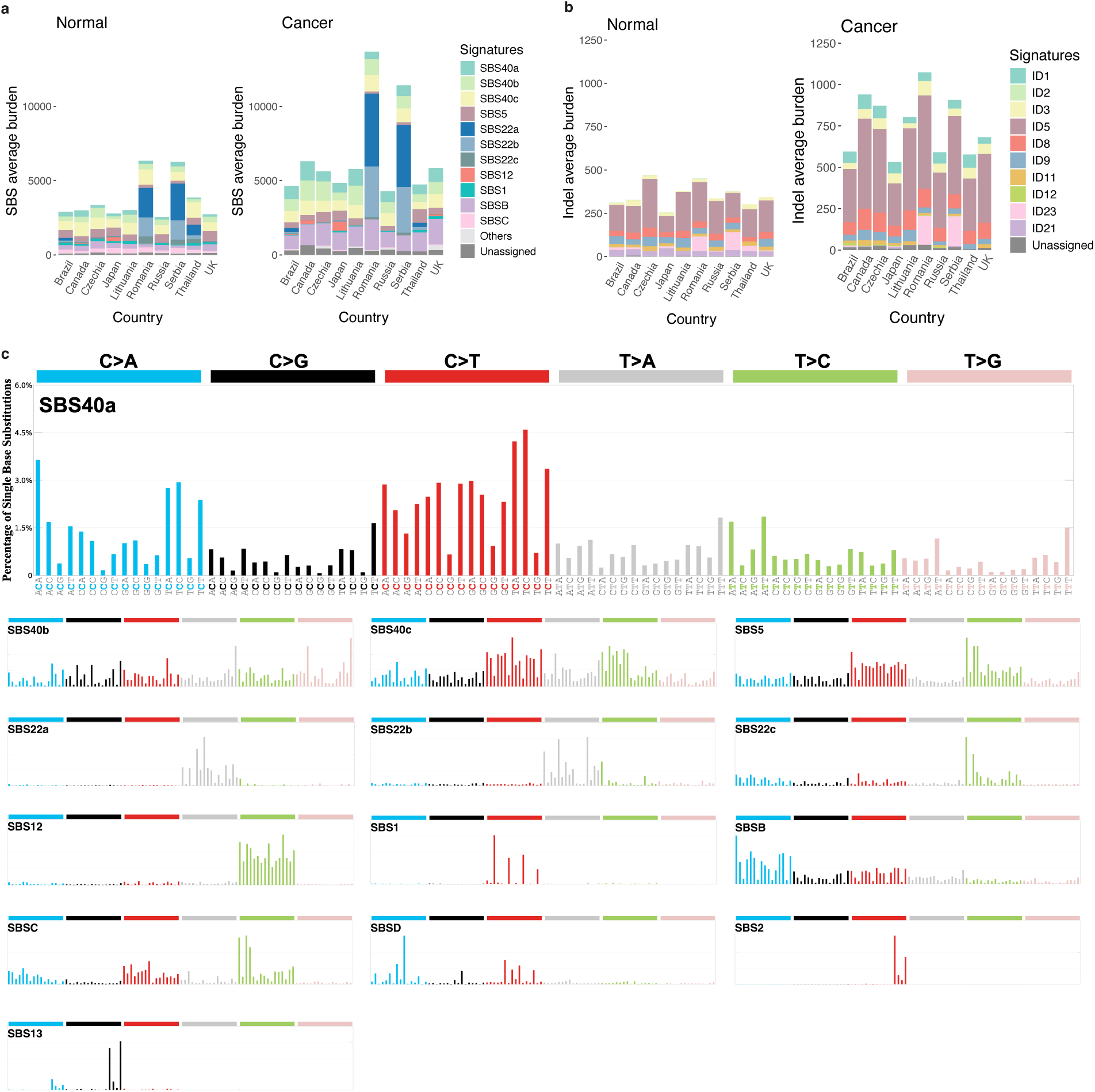
Mutational signatures in normal kidneys. **a**, Average attribution for SBS mutational signatures across countries in 299 bulk normal kidney cortex samples (left) and 291 matched cancers (right). Normal samples are from Brazil (*n* = 40), Canada (*n* = 7), Czechia (*n* = 67), Japan (*n* = 64), Lithuania (*n* = 11), Romania (*n* = 33), Russia (*n* =37), Serbia (*n* = 12), Thailand (*n* = 4), UK (*n* = 24). Matched cancer genomes are available for all donors except one from Czechia, one from Japan, one from Romania, and four from the UK. One cancer sample from Serbia (PD47592a) was excluded due to hypermutation caused by loss of *XPC*. Others: SBS2, SBS13, SBS18, SBSD. Unassigned: signatures contributing to <5% of total burdens in each sample. **b**, Average attribution for indel mutational signatures across countries in 299 bulk normal kidney cortex samples (left) and 290 matched cancers (right). One additional UK cancer case (PD50441a) with indel hypermutation dominated by ID2 was excluded. Unassigned: signatures contributing to <5% of total burdens in each sample. **c**, SBS mutational signatures in normal kidneys.

SBSA correlates closely with SBS22a and SBS22b caused by aristolochic acid exposure^18^, appears to be a further separating component of the composite aristolochic acid signature and has, therefore, been designated SBS22c. SBSD was present in a single normal kidney sample and is similar to a signature found in platinum chemotherapy-exposed liver^19^, although this individual has no history of chemotherapy. SBSB and SBSC are discussed below.

All indel signatures from *de novo* extraction were decomposed into COSMIC reference signatures: clock-like ID5, tobacco smoking-associated ID3, aristolochic acid-associated ID23, and signatures with unknown etiologies including ID8, ID9, ID11 and ID21^7,12,20^(Extended Data Fig.4). Due to the scarcity of double-base substitutions (DBS) given the duplex coverage, *de novo* DBS signatures were not extracted and DBS mutations were instead attributed to the signatures reported in ccRCC^12^: DBS2, DBS4, DBS9, DBS20 and DBS_C. DBS2 burden was confirmed to be associated with smoking history (*P* = 4.7 × 10^−5^).

SBS1 (due to deamination of 5-methyl cytosine), SBS5 (of unknown mechanism) and SBS40a (of unknown mechanism) are present in most normal cell types, accrue at a more-or-less linear rate with age throughout life, and are thought to be due to endogenous mutational processes^21,22^. SBS2 and SBS13 are caused by activity of members of the APOBEC family of cytidine deaminases^23–25^, were found in three bulk kidney normal samples and have been reported in other normal cell types including small intestine^26^, urinary bladder^27^ and bronchial epithelia^10^.

SBS22a, SBS22b and SBS22c are caused by exogenous, lifestyle or environmental, exposure to aristolochic acids, renal toxins and mutagenic carcinogens produced by species of the *Aristolochia* plant genus^18^. These signatures show transcriptional strand bias of mutations (Extended Data Fig.5a), a feature typically associated with exogenous mutagens which form bulky DNA adducts that distort the double helix and recruit transcription-coupled nucleotide excision repair to remove them^13^. Normal bulk kidney samples with these signatures clustered in Romania and Serbia (Fig.2a and Extended Data Fig.6a–c), as previously seen in ccRCC^12^, indicating an epicentre of intense local aristolochic acid exposure in Southeast Europe the reason for which remains unclear. The SBS22a, SBS22b and SBS22c mutation loads account for the high overall mutation burdens in these countries and are consistent with aristolochic acids being ingested, absorbed, and systemically circulated in the blood, thereby exposing normal internal body tissues.

SBS12 is found, commonly and almost exclusively, in ccRCC^7,12^ and hepatocellular carcinomas^7,28^ from Japan. This distinctive geographic distribution and the presence of transcriptional strand bias prompted the hypothesis that SBS12 is caused by regional exposure to a currently unknown exogenous mutagenic agent. The observation here of SBS12 exclusively in normal kidney cells of individuals from Japan (Fig.2a and Extended Data Fig.6d) lends important support to the hypothesis of exogenous origin and excludes the possibility that it is due to a mutational process recruited during clonal evolution of the cancers in which it is found.

SBS40b has only been reported in ccRCC^12^. It was detected in ccRCC from all 11 countries previously studied but showed geographic variation in prevalence and mutation burden which positively correlated with ccRCC national incidence rates, age, male sex, and indices of impaired kidney function^12^. In bulk normal kidney cortex, SBS40b burden is positively associated with age (adjusted *P* = 5.9 × 10^−8^) and male sex (adjusted *P* = 0.009), showing modest transcriptional bias (Extended Data Fig.5b), and its mutation rate correlates significantly with age-standardized ccRCC incidence rates (Spearman’s ρ = 0.84, adjusted *P* = 9.7 × 10^−3^). The geographic variation in SBS40b burden found in ccRCC was confirmed in normal kidney samples, with Czechia showing a high burden (267 more mutations, 95% CI: 178-355, adjusted *P* = 2.7 × 10^−7^) and Japan a low burden (236 less mutations, 95% CI: 144-328, adjusted *P* = 1.2 × 10^−5^) compared to the average over all countries (Fig.2a and Extended Data Fig.7). The presence of SBS40b in normal kidney, and its geographically variable mutation burden, raise the possibility that it is also due to a systemically circulating mutagen of exogenous origin. If so, the agent would be ubiquitous and may, similarly to aristolochic acids, be both a renal mutagen and toxin. Alternatively, however, SBS40b could be due to a mutational process of endogenous origin which is confined to the kidney, perhaps, consequent on kidney damage caused by factors which show geographical variation.

SBS40c has only been reported in ccRCC without evidence of geographic variation in prevalence^12^. It shows transcriptional strand bias (Extended Data Fig.5a). Its presence in normal kidney cells excludes its origin as a mutational process operating during neoplastic transformation and raises the possibility that it is also generated by a systemically circulating mutagen. However, as for SBS40b, other explanations cannot be excluded.

SBSB is similar to SBS4 (0.85 cosine similarity), which is caused by exposure to tobacco smoke chemicals^29^, and exhibits similar transcriptional strand bias. SBSB was previously extracted by others from kidney cancers^30^ and has been associated with tobacco smoking in lung cancer^31,32^. In our studies, its presence in normal kidney and in ccRCC correlates with a smoking history (*P* = 1.3 × 10^−4^ and *P* = 4.5 × 10^−5^ respectively) although it was present in two-thirds of normal kidney samples, some from non-smokers. Therefore, SBSB is plausibly, at least in part, due to chemicals present in tobacco smoke, and potentially other exposures, which are absorbed and systemically circulated by the blood to the normal kidney. Its differences from SBS4 may reflect that only a subset of the chemicals to which smokers’ bronchial epithelial cells are directly exposed reach the kidney, that they are processed by different metabolic pathways in different tissues, or that SBSB is a composite of multiple signatures with similar profiles. SBSC was not previously reported in ccRCC and is characterized predominantly by T>C substitutions at A<u>T</u>A and A<u>T</u>T trinucleotides.

### Variation in mutation rates and mutational signatures across cell types

To investigate differences in mutation rates and signatures between different kidney cell types, microdissected glomeruli, proximal tubules, medulla and distal tubules from 84 individuals were sequenced for somatic mutations using NanoSeq. It should be noted that, to varying extents, all these structures are cell type mixtures and this will be reflected in the somatic mutations reported by NanoSeq sequencing.

Samples from 15 individuals from the UK, aged 9–85 years, without a history of renal cancer, or known region-specific exposure, or severe chronic kidney disease were used to estimate mutation rates in different parts of the nephron in normal kidney from healthy people. In these individuals, proximal tubule cells accumulated 60 SBS (95% CI: 52–68) and 7.8 indels (95% CI: 5.9–9.6) per genome per year. Excluding skin epidermis with ultraviolet light exposure^9^ and lung bronchial epithelial cells with tobacco smoke exposure^10^, these mutation rates exceed or are comparable to the highest observed in normal human tissues analyzed thus far despite the low mitotic division rate of proximal tubule cells^33^ and the absence of a known direct exogenous mutagen exposure (Extended Data Table 1. Examples of actively dividing tissues include colorectal epithelium, which accumulates ∼44 SBS and ∼1–2 indels per genome per year, and small intestinal epithelium, which accumulates ∼42–51 SBS and ∼2–4 indels per genome per year. Other normal tissues generally acquire 15–40 SBS per genome per year^14,22,26,27,34–43^.) Other nephron structures exhibited lower mutation rates: glomerulus 38 SBS (95% CI: 34–43) and 2.0 indels (95% CI: 1.7– 2.3); medulla 42 SBS (95% CI: 31–53) and 3.9 Indels (95% CI: 1.6–6.3); distal tubule 24 SBS (95% CI: 20–27) and 2.0 Indels (95% CI: 1.7–2.2).

Comparison of mutational signatures between the different nephron structures across 84 individuals revealed striking cell type-specific mutational patterns. Proximal tubules showed similar or lower mutation burdens of SBS1, SBS5 and SBS40a, signatures of likely endogenous origin, to other nephron components from the same individuals. By contrast, in individuals known to have experienced aristolochic acid exposure through the presence of SBS22a, SBS22b and SBS22c in ccRCC and normal kidney, the mutation burdens of SBS22a, SBS22b, SBS22c and ID23 (the indel signature caused by aristolochic acid exposure) were substantially higher in proximal tubules than in glomeruli, medulla, and distal tubules (Fig.3d and Fig.4a. SBS22a: 470 more mutations, 95% CI: 246–694, adjusted *P* = 1.7 × 10^−4^; SBS22b: 631 more mutations, 95% CI: 368–894, adjusted *P* = 1.5 × 10^−5^; SBS22c: 52 more mutations, 95% CI: 11–92, adjusted *P* = 0.025; ID23: 26 more mutations, 95% CI: 11–41, adjusted *P* = 1.4 × 10^−3^). Indeed, the real differences may be even greater because laser-capture microdissection cannot completely prevent contamination by other cell types. These differences in aristolochic acid-induced mutation burdens may reflect normal functions of proximal tubules, including reabsorption of organic compounds from the glomerular filtrate and secretion from the blood into the urine^44,45^. It is conceivable that in the process of active absorption and transportation of aristolochic acids, proximal tubule cells generate high intracellular concentrations which result in elevated levels of mutagenesis. However, other explanations cannot be excluded.

**Fig. 3.**
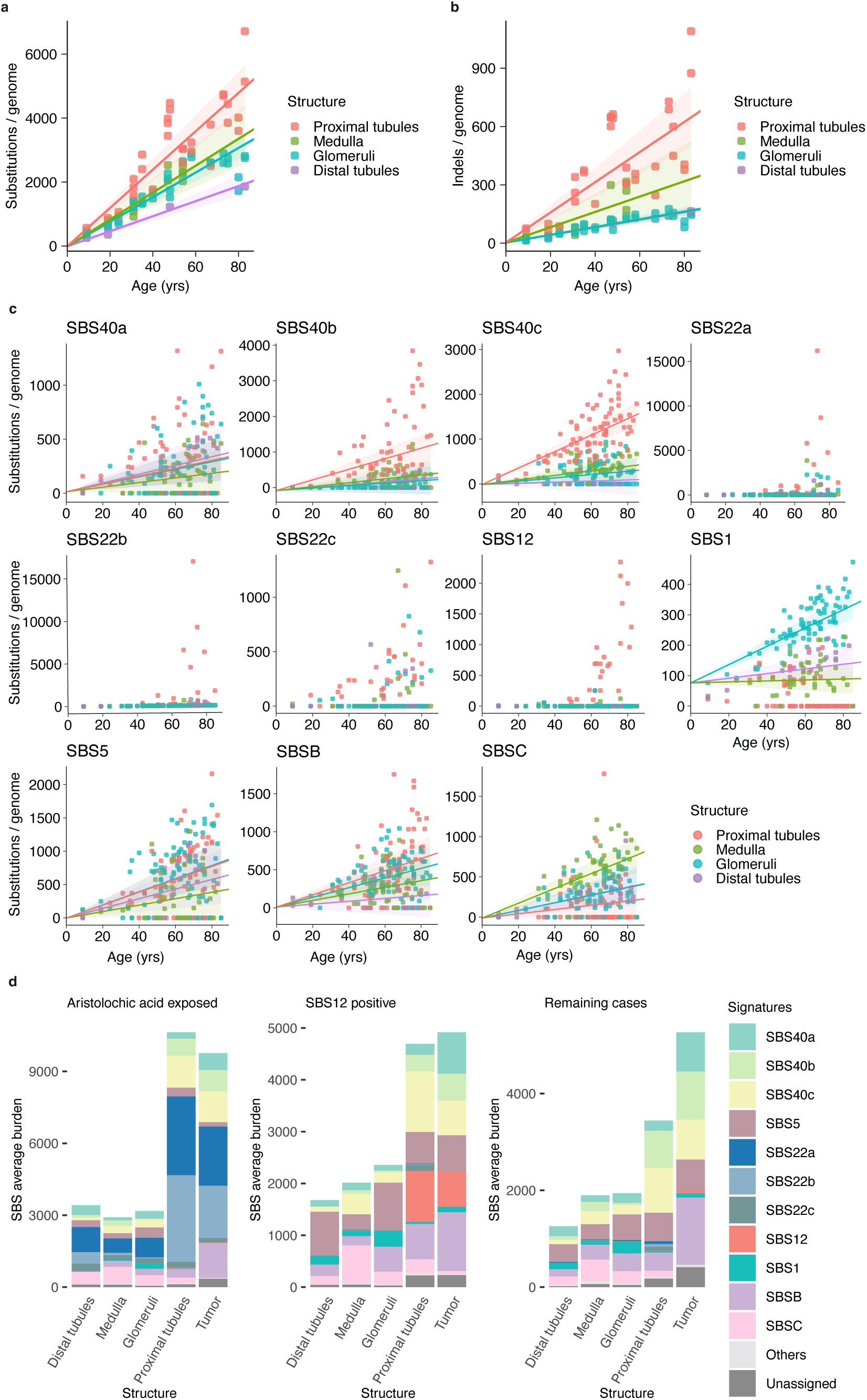
Mutation rates and burdens of mutational signatures across different kidney structures. **a,** SBS burden versus age, showing regression lines for proximal tubules, glomeruli, distal tubules and medulla. Regression lines were estimated using linear mixed-effects models. Error bands represent 95% confidence intervals for the fixed effects of age. *N* = 83 libraries from 15 individuals without cancer or other diseases. **b**, Indel burden versus age, showing regression lines for proximal tubules, glomeruli, distal tubules and medulla. Regression lines were estimated using linear mixed-effects models. Error bands represent 95% confidence intervals for the fixed effects of age. *N* = 83 libraires from 15 individuals without cancer or other diseases. **c**, Mutational burden versus age for every signature with >1% total burden contribution. Regression lines were estimated using linear mixed-effects models. Error bands represent 95% confidence intervals for the fixed effects of age. *N* = 468 libraries from 84 individuals, libraries from the same structure and same individual were pooled (*N* = 238) and shown as average. **d**, Average attribution for SBS signatures across different nephron structures from ccRCC patients and corresponding cancer genomes. Left, aristolochic-acid exposed cases (with >5% contributions from SBS22a, SBS22b and SBS22c in their cancer genomes, 14 donors). Middle, SBS12-exposed cases (with >5% contributions from SBS12 in their cancer genomes, 19 donors). Right, all remaining cases (35 donors). Others: SBS2, SBS13, SBS18, SBSD. Unassigned: signatures contributing to <5% of total burdens in each sample.

SBS12, SBS40b, and SBS40c showed a similar pattern of higher mutation burdens in proximal tubules compared to other nephron components (Fig.3d and Fig.4b–d. SBS12: 163 more mutations, 95% CI: 116–211, adjusted *P* = 5.6 × 10^−10^; SBS40b: 431 more mutations, 95% CI: 348–514, adjusted *P* = 7.3 × 10^−21^; SBS40c: 658 more mutations, 95% CI: 572–744, adjusted *P* = 8.5 × 10^−39^). SBSB and ID3, which at least in part are likely due to tobacco smoke chemicals, were moderately enriched in proximal tubules (SBSB: 136 more mutations, 95% CI: 76–195, adjusted *P* = 4.0 × 10^−5^; ID3: 11 more mutations, 95% CI: 4–17, adjusted *P* = 0.002). Thus, signatures due to known systemically circulating mutagen exposures and many signatures of uncertain origin showed higher mutation burdens in proximal tubules than in other nephron structures. Signatures showing geographic variation in burden in bulk normal kidney cortex exhibited the same pattern even more prominently in proximal tubules: SBS12 contributed 536 more mutations in Japan than the mean of all countries (95% CI: 312–759, adjusted *P* = 2.6 × 10^−4^), and SBS40b contributed to 737 fewer mutations in Japan (95% CI: 398–1076, adjusted *P* = 0.001) and 1062 more mutations in Czechia (95% CI: 573–1551, adjusted *P* = 0.001). These observations add weight to the previously hypothesized origin of SBS12 in Japan as an exogenous exposure and further highlight the possibility of this origin for SBS40b and SBS40c. SBSC was present in all nephron structures but with the highest mutation rate in the medulla; SBS1 showed a higher mutation rate in glomeruli than in other nephron structures (Fig.3c–d).

**Fig. 4.**
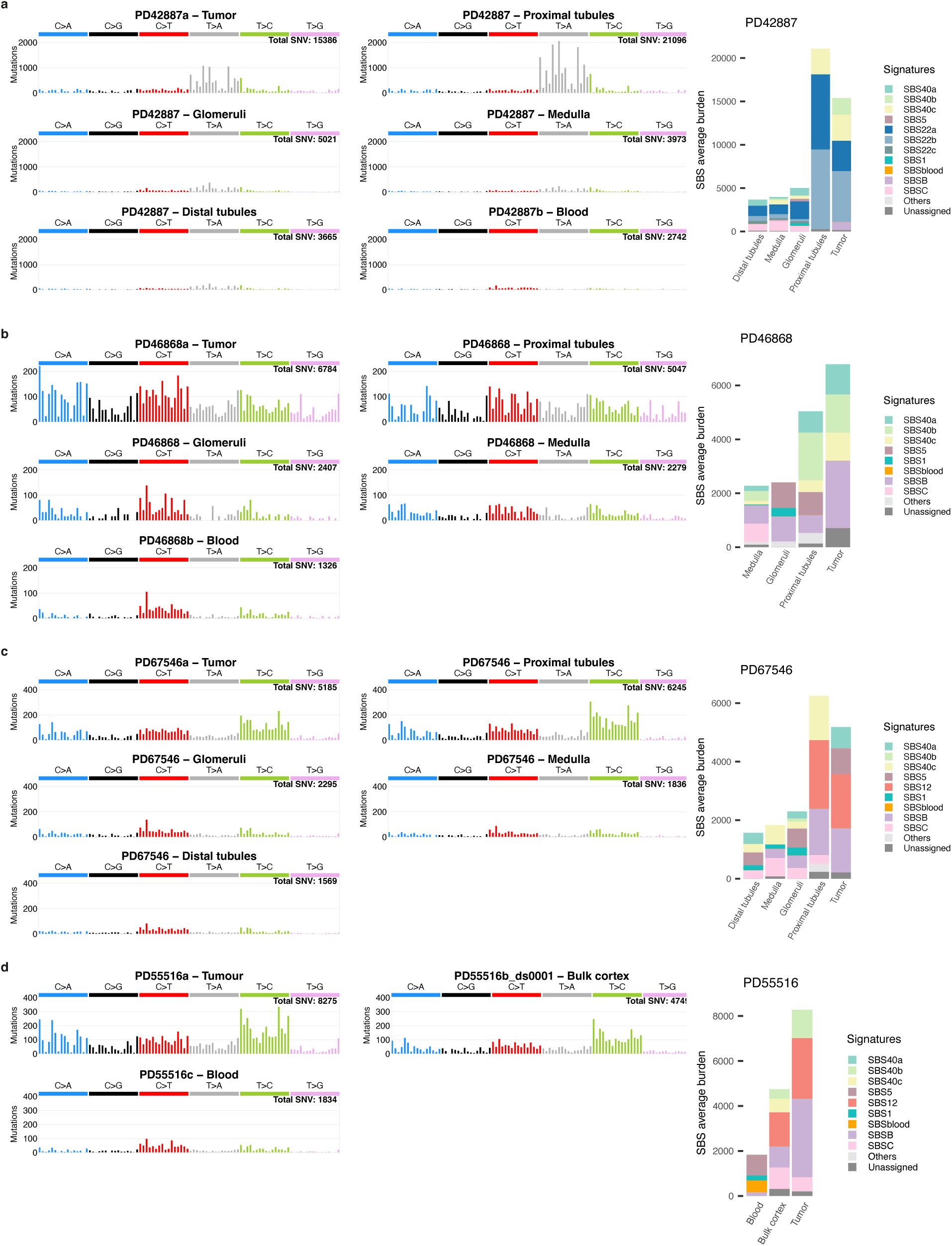
Examples of cell-type specific mutational signatures. **a**, SBS mutational spectra and signature attribution in PD42778, showing high levels of SBS22a and SBS22b in cancer and proximal tubules. Others: SBS2, SBS12, SBS13, SBS18, SBSD. Unassigned: signatures contributing to <5% of total burdens in each sample. **b**, SBS mutational spectra and signature attribution in PD46868, showing high levels of SBS40b and SBS40c in cancer and proximal tubules. Others: SBS2, SBS12, SBS13, SBS18, SBS22a, SBS22b, SBS22c, SBSD. Unassigned: signatures contributing to <5% of total burdens in each sample. **c**, SBS mutational spectra and signature attribution in PD67546, showing high levels of SBS12 in cancer and proximal tubules. Others: SBS2, SBS13, SBS18, SBS22a, SBS22b, SBS22c, SBSD. Unassigned: signatures contributing to <5% of total burdens in each sample. **d**, SBS mutational spectra and signature attribution in PD55516, showing SBS12 was absent in blood while present in cancer and normal kidney cortex. Others: SBS2, SBS13, SBS18, SBS22a, SBS22b, SBS22c, SBSD. Unassigned: signatures contributing to <5% of total burdens in each sample.

Elevated indel burdens in proximal tubules were predominantly explained by increased ID5, ID8, ID11, ID9 (and ID3, see above) compared to other nephron structures (Extended Data Fig.4. ID5: 187 more mutations, 95% CI: 158–216, adjusted *P* = 2.4 × 10^−29^; ID8: 32 more mutations, 95% CI: 26–38, adjusted *P* = 1.2 × 10^−20^; ID11: 27 more mutations, 95% CI: 23–32, adjusted *P* = 7.7 × 10^−29^; ID9: 18 more mutations, 95% CI: 12–24, adjusted *P* = 3.7 × 10^−7^;).

NanoSeq sequencing of normal peripheral blood samples from 272 individuals for somatic mutations did not reveal the geographic variation in mutation burden observed in ccRCC or normal kidney tissue. Mutational signature analysis revealed, as expected, primarily SBS1, SBS5 and SBSblood^42^. However, there was no evidence of aristolochic acid induced SBS22a, SBS22b and SBS22c in individuals with these signatures in their ccRCC and normal kidney samples, even when mutation burdens were high (Fig.4). Similarly, SBS12, SBSB, and SBS40b were not observed in blood from individuals showing them in normal kidney and ccRCC.

### Normal kidney cell and renal clear cell cancer genomes

Many thousands of whole cancer genome sequences have been generated over the last two decades^46^, but the somatic mutation landscapes of normal cells from which a cancer has developed with the cancer from the same person have rarely been directly compared due to technical obstacles previously preventing the detection of somatic mutations in normal tissues.

The SBS and indel mutation burdens of ccRCC were generally higher than those of normal kidney cells from the same individual. The differences were, however, modest with a median increase of 30% SBS and 37% indels for proximal tubules, although a small number of ccRCCs exhibited substantial increases compared to their normal kidney tissues. Mutation burdens attributable to SBS22a, SBS22b, and ID23 due to aristolochic acid exposure were similar or slightly higher in proximal tubules compared to ccRCC (Fig. 3d, Fig. 4a and Extended Data Fig.4b). Their mutation burdens were lower in bulk normal kidney than in ccRCC (Fig. 2a–b) probably because proximal tubule cells, from which ccRCC derives, constitute only a subset of cells in the bulk normal kidney samples. Similar patterns were observed for SBS12, SBS40b and SBS40c. Thus, exposures to circulating mutagens increase mutation burdens primarily in normal cells, with limited impact during later clonal evolution of the cancer.

ccRCC exhibited three SBS signatures not found in normal kidney samples from the same individuals: SBS18 (due to DNA damage by reactive oxygen species)^47^, SBS21 and SBS44 (due to defective DNA mismatch repair^48,49^ and SBS_H. SBS_H is likely due to biallelic loss of *XPC*, which encodes a component of the nucleotide excision repair machinery^50,51^, and was found in one individual with a germline *XPC* truncating mutation and somatic loss of the wild type *XPC* allele in the kidney cancer. The higher total single-base substitution mutation burdens of ccRCC compared to proximal tubule cells from the same individuals were, however, largely attributable to additional burdens of SBS40a and SBSB (Fig. 3d). SBSC, which is predominantly present in normal medulla, was not detected in ccRCC. Indel signatures ID1 and ID2 (due to polymerase slippage during DNA replication), as well as ID12 of unknown etiology were not extracted in normal kidneys. These signatures, therefore, represent mutational processes which are more active during neoplastic clonal evolution than in normal kidney tissue.

## Discussion

Exogenous mutagen exposures are expected to cause mutations in normal tissues. This study provides the foundations for the interpretation of mutational epidemiology surveys conducted through kidney tissues. The results show that somatic mutagenesis of normal human kidney cell genomes is complex, involving a substantial number of mutational processes, and that mutation burdens are higher in proximal tubule cells than in most other normal tissues. Some signatures are due to known exogenous environmental and lifestyle mutagen exposures and others to known endogenous mutational processes. The remainder are of uncertain origin. Ultimately, clarification of their origins will depend on improved mechanistic understanding which, for exogenous exposures, will include epidemiological studies and experimental approaches to identify the causative mutagens.

In the interim, however, the results highlight features distinguishing mutational signatures due to exogenous exposures, even when the causative agents are unknown. First, presence of the mutational signature in normal as well as cancer cells helps to differentiate them from mutational signatures of endogenous processes recruited during neoplastic clonal evolution. Previously technologically challenging, this can now be determined in all normal tissues using NanoSeq and other methods. Second, exogenous exposures often cause variation in mutation burden of the mutational signature which is not attributable to age and which can be between populations or geographic regions, between individuals within a population, or over time. Such variation due to exogenous exposures can be large but lesser degrees of variation could, in principle, arise from more ubiquitous exogenous exposures. Nevertheless, alternative explanations of such differences should be considered, including inherited genome variation or exogenous factors influencing the mutation rates of endogenous mutational processes. Third, intrinsic features of the mutational signature can help inform on exogenous exposures. Transcriptional strand bias is observed in mutational signatures induced by most known exogenous mutagens (e.g. SBS4, SBS7, SBS22, SBS24, and many chemotherapies). Nevertheless, it is occasionally observed in some endogenous mutational signatures found in cancers due to acquired defects in DNA repair (e.g. SBS8, SBS26) and it is also conceivable that some exogenous mutagens do not induce it. Fourth, high mutation burdens are present in certain normal cell types. Renal proximal tubule cells exhibit particularly high SBS22a and SBS22b mutation burdens due to aristolochic acids presumably due to their intrinsic biological properties. The similar pattern observed for SBS12 in samples from Japan suggests that this may apply more widely to exogenous mutagens. Other cell types may have similar capabilities and the common presence of SBS22a and SBS22b in hepatocellular carcinomas^18,52^, together with SBS12 in Japanese cases^7,28^, suggests that hepatocytes constitute a further example. However, some endogenous mutational signatures also exhibit high mutation burdens in particular cell types without known exogenous exposures and this needs to be retained in consideration. None of these features is, therefore, sufficient or necessary to establish an exogenous mutagen as the cause of a mutational signature and there exist exceptions for each. Together, however, they form a composite profile of mutational signatures caused by exogenous mutagens which constitutes a foundation for future elaboration and consolidation. This framework strengthens the candidature of SBS12, and highlights that of SBS40b and SBS40c, as outcomes of exogenous mutagen exposures.

In principle, the ability of exogenous mutagens to generate specific mutational signatures in human tissues provides an opportunity to survey lifetime exposures in general populations, at scale and internationally. Samples of reporter cell types for this purpose should, ideally, be relatively easily obtained, for example buccal epithelium or blood. However, peripheral blood does not appear to report SBS22a and SBS22b in aristolochic acid-exposed individuals, at least at the exposure intensities surveyed. Therefore, the remarkable propensity of kidney proximal tubule cells (and the ccRCCs which arise from them) to report systemically circulating mutagens of exogenous origins offers an alternative approach to conducting future surveys of this nature.

## Materials and Methods

### Ethics and Materials

The 303 donors with ccRCC were recruited in the participating countries through an international network coordinated by the International Agency for Research on Cancer (IARC/WHO). Among them, 267 donors had their ccRCC whole-genome sequencing previously published, and 29 donors had their ccRCC whole-gnome sequenced additionally in this study. The normal kidney samples were procured during nephrectomy for kidney tumor removal, and were sectioned, H&E stained, reviewed by a histopathologist with expertise in kidney disease diagnosis to ensure they are normal, with no obvious histopathological abnormality. Ethical approvals were first obtained from each Local Research Ethics Committee and Federal Ethics Committee when applicable, as well as from the IARC Ethics Committee.

The second cohort of 16 UK donors without kidney cancer included samples from autopsy of a child died with high-grade midline gliomas (PD50297), approved by NRES Committee East Midlands—Derby with REC reference 08/H0405/22+5; samples from AMSBio (commercial supplier)(PD43850, PD43851, PD56385 and PD56386), whose donors had died of causes not related to cancer, approved by the London-Surrey Research Ethics Committee with REC reference 17/LO/1801); and the remainder being cadaveric kidneys that were collected for kidney transplantation but subsequently declined (approved by NRES Committee East of England—Cambridge East with REC reference 12/EE/0202 and 16/EE/0014).

### Bulk and low-input DNA extraction

Extraction of DNA from fresh frozen bulk normal kidney cortex and periphery blood was centrally conducted at IARC/WHO, with exception for 30 Japanese UK normal kidney cases which were extracted at the local center following a similarly procedure. These samples were examined and selected by pathologist to have minimal or no medulla component. For samples being micro-dissected, a standard protocol established at Wellcome Sanger Institute for low-input DNA extraction was adapted. Fresh frozen biopsies were fixed, embedded, sectioned and stained before library preparation. PAXgene FIX Kit (PreAnalytiX, 765312) was used for fixation. Subsequent paraffin embedding was applied for higher quality morphology of the tissue. Biopsies were sectioned to 10 µm, fixed to 4 µm PEN membrane slides (11600288, Leica) and stained with hematoxylin and eosin. Different structures were isolated using laser-capture microscopy (LMD7000, Leica) and collected in Eppendorf LoBind® PCR 96-well plates. Collected samples were lysed using ARCTURUS PicoPure DNA extraction kit (Applied Biosystems) according to the manufacturer’s instructions. For normal kidney samples received without DNA extract, bulk cut of kidney cortex from the micro-dissected biopsy were collected in Eppendorf DNA LoBind® tubes and extracted DNA using Qiagen micro-DNA extraction kit (catalogue number 56304) according to the manufacturer’s instructions.

### NanoSeq library preparation and sequencing

NanoSeq libraries were prepared following the standard protocol established at Wellcome Sanger Institute ^14^: genomic DNA or lysed tissue microbiopsies were purified using 100 μl of a 50:50 water:AMPure XP bead mixture and eluted in 20 μl nuclease-free water (NFW). Then, the bead suspension was taken forward into an on-bead fragmentation reaction in a final volume of 25 μl including 2.5 μl 10× CutSmart buffer (500 mM potassium acetate, 200 mM Tris-acetate, 100 mM magnesium acetate, 1 mg/ml BSA, pH 7.9 at 25 °C), 0.5 μl 5U/μl HpyCH4V and 2 μl NFW. Fragmentation reactions were incubated at 37 °C for 15 min, purified with 2.5× AMPure XP beads and resuspended in 15 μl NFW. Fragmented DNA was A-tailed in incubated at 37 °C for 30 min before ligation of Duplex Seq Adapters (IDT, 1080799). Reactions were incubated at 20 °C for 20 min and subsequently purified with 1× AMPure XP beads and resuspended in 50 μl of NFW. The choices of restriction enzyme (HpyCH4V) and fragment size selection (250–500 bp) restrict the coverage to around 30% of the human genome.

From diluted NanoSeq libraries (0.1-0.6 fmol DNA in 25 μl), paired-end sequencing reads (150 bp) were generated using Illumina NovaSeq 6000 platform. Sequences were aligned to the human reference genome (GRCh37) using BWA-MEM^53^, using the -C option to append barcode sequences to alignments. Alignments were sorted by coordinate and read duplicates were marked using bamsormadup^54^. Reads were filtered when they were marked as unpaired, optical duplicate, supplementary, QC fail, unmapped or secondary alignments. A median of 3 Gb effective duplex coverage was achieved for each library.

### Variant calling from NanoSeq data

For both single-base substitutions and indels, standard 20× whole-genome sequencing data of blood or bulk normal kidney from the same individuals was used as matched normal to exclude germline mutations. Single-base substitution calling was done by an in-house NanoSeq analysis pipeline (https://github.com/cancerit/NanoSeq) with a set of customized filters^14^, with genomic mask for common SNP and TGCA restriction site artefact masking: (1) each read bundle should have at least two duplicate reads from each original strand to support a mutation. (2) The consensus base quality score should be at least 60. (3) The minimum difference between the primary alignment score (AS) and the secondary alignment score (XS) should be higher than 50. (4) The average number of mismatches in a group of reads (forward or reverse) should not be higher than 2. (5) No 5 clips are allowed. (6) No improper pairs are allowed in the read bundle. (7) Base calls in within 8 bp from the 5 or 3 ends are discarded. (8) Reads in the read bundle must contain no indels. (9) The matched normal must have over 15× coverage at the given site. (10) Mutations seen with a frequency higher than 0.01 in the matched normal are excluded. (11) Sites overlapping the common SNP and noisy sites are masked.

Indel calling was done by bcftools^55^ integrated in the same in-house NanoSeq analysis pipeline, with a set of customized filters^14^: (1) the indel should be contained in at least 90% of forward and reverse reads of the read bundle. (2) The minimum difference between the primary alignment score (AS) and the secondary alignment score (XS) should be higher than 50. (3) No 5′ clips are allowed. (4) The matched normal must have over 16× coverage at the given site. (5) Indels close to read ends (10 bp) were not called. (6) Sites overlapping the common SNP and noisy sites are masked. (7) The proportion of indels in the matched normal within the ±5-bp window around the candidate somatic indel should not be higher than 1%.

### Detection of contamination in NanoSeq samples

Because NanoSeq detects mutations at low frequency, it is very sensitive to DNA contamination from other individuals. The in-house NanoSeq analysis pipeline integrated VerifyBamID2^56^ to estimate the extent of contamination in NanoSeq samples. This method leverages individual-specific allele frequencies projected from reference genotypes onto principal component coordinates to estimate DNA contamination, and does not require prior knowledge about the genetic ancestry of the contaminating sample. Samples with > 0.5% possible contamination were excluded from downstream analysis.

### Correction of NanoSeq mutation matrix

The number of substitutions observed in each trinucleotide context is biased by the masks (higher filtering of CpG sites), the restriction sites (trinucleotides overlapping TGCA being depleted) and the background trinucleotide frequency of ∼30% genome covered by NanoSeq. To adjust for this bias, a ratio of genome-wide trinucleotide frequency to experimental trinucleotide frequency was calculated. The observed substitution burden within each trinucleotide context was multiple by this ratio of the corresponding trinucleotide context to obtain a trinucleotide context-adjusted substitution burden.

In order to make fair comparison between NanoSeq mutation burden and that from standard whole-genome sequencing, the mutation burden per normal diploid human genome were estimated by trinucleotide frequency corrected NanoSeq burden multiplying the ratio of the entire diploid GRCh37 reference genome size to the total number of duplex sequenced bases.

### Whole-genome sequencing and variant calling

Whole-genome sequencing was performed on the Illumina NovaSeq 6000 platform with target coverage of 40× for tumors and 20× for matched normal tissues. Variant calling was performed using the standard Sanger bioinformatics analysis pipeline (https://github.com/cancerit). Copy number profiles were determined first using the algorithms ASCAT^57^ and BATTENBERG^58^. Single nucleotide variants (SNVs) were called with cgpCaVEMan^59^ using the copy number profile and purity values determined from ASCAT, indels were called with cgpPINDEL^60^. A second variant caller, Strelka2^61^, was run for SNVs and indels as consensus variant calling to eliminate algorithm specific artefacts. Additional filters on ASMD (median alignment score of reads showing the variant allele) and CLPM (median number of soft-clipped bases in variant supporting reads) (ASMD ≥ 140 and CLPM = 0) were applied to remove potential false positive calls.

### Mutational signature extraction and decomposition

Mutational signatures in normal kidneys were extracted from all bulk normal cortex samples and LCM biopsies. To minimize noises in signature extraction resulting from the low-input DNA sequencing for LCM biopsies, if multiple biopsies from the same structure in the same individual were available, their raw mutation counts in each mutation category were merged for mutational signature extraction. After merging, those with at least 50 mutations were kept for mutational signature extraction. Two independent algorithms were used to extract mutational signatures from single-base substitution and indel matrices: (1) mutational signature extraction was run on each mutational matrix using HDP^15^ (v0.1.6, https://github.com/NickWilliamsSanger/hdp) without priors and hierarchy, in 20 independent chains for 150,000 iterations and with a burn-in of 50,000. (2) *De novo* mutational signatures were extracted from each mutational matrix using SigProfilerExtractor^16^ (v1.1.24, https://github.com/AlexandrovLab/SigProfilerExtractor) with NMF_init = nndsvd_min, maximum_signatures = 20 (substitutions) or 10 (indels), nmf_replicates=500.

*De novo* signatures from normal kidneys were then decomposed using SigProfilerAssignment^62^ (v0.1.4, https://github.com/AlexandrovLab/SigProfilerAssignment) to a known list of COSMIC reference signatures in kidney cancer (SBS40a, SBS40b, SBS40c, SBS22a, SBS22b, SBS5, SBS12, SBS18, SBS1, SBS2, SBS13, SBS21, SBS44, SBS4, ID1, ID2, ID3, ID5, ID8, ID9, ID11, ID12, ID23)^12^. *De novo* signatures that cannot be reconstructed with cosine similarity over 0.95 for single-base substitution signatures or over 0.85 for indel signatures were decomposed against all COSMIC v3.4 reference signatures^17^, and those still cannot be reconstructed by the previous cosine similarity standard were defined as novel signatures.

The final signatures were supported by one or more of the following criteria: (1) signatures reported by both algorithms, (2) signatures clearly supported by individual mutational spectra, (3) signatures associated with specific sample characteristics (such as tissue, country of origin), and (4) COSMIC reference signatures decomposed from the extracted signatures. The final list consists of the decomposed COSMIC reference signatures and the novel signatures, with the novel signature SBSB replacing reference signature SBS4, as SBS4 reported in kidney cancer originally came from the decomposition of a version of this signature extracted in kidney cancer. For two noisy novel signatures, in the final version, more data (962 available kidney cancer whole-genome data from a previous study^12^ and 29 sequenced in this study) were added into the *de novo* signature extraction for a clearer version (SBSB), or LCM data were removed to further reduce noises (SBS22c). For blood, signatures from *de novo* extraction were decomposed to normal kidney signatures plus additional previously reported blood signatures (SBSblood, SBS7a, SBS8, SBS9, SBS17b, SBS19)^42,63^.

### Mutational signature attribution

The final list of signatures present in normal kidney, along with additional COSMIC reference signatures reported in kidney cancer, were attributed for each sample using the MSA signature attribution tool (v2.2, https://gitlab.com/s.senkin/MSA)^64^. MSA applies an optimized penalty (params.no_CI_for_penalties = FALSE) to discourage the inclusion of mutational signatures that do not substantially improve the reconstruction of the input mutational profile. For a given sample, mutational signatures are attributed by non-negative least squares (NNLS) using a candidate set of signatures, and penalties are applied to reduce overfitting by removing signatures that do not meaningfully increase the similarity between the observed and reconstructed profiles. Penalty values were pre-optimized using simulated data, such that the specificity for the detection of each signature exceeded 95%. Signatures contributing to less than 5% mutations in each sample were grouped into “unassigned” component. Normal kidney and kidney cancer signatures (SBS40a, SBS40b, SBS40c, SBS22a, SBS22b, SBS22c, SBS5, SBS12, SBS18, SBS1, SBS2, SBS13, SBS21, SBS44, SBSB, SBSC, SBSD, ID2, ID3, ID5, ID8, ID9, ID11, ID12, ID21, ID23) were attributed to all normal kidney and cancer samples in one combined attribution. DBS signatures were not attributed microdissected samples because of limited raw DBS mutations. Signatures extracted and decomposed from blood samples (SBS1, SBS5, SBSB, SBSblood) were attributed to blood.

### Mutation rate in normal kidney

Linear mixed-effects models were implemented using the nlme package (v3.1.168) to estimate structure-specific mutation rates for single-base substitution (SBS) and indel mutation burdens.

To estimate the total mutation rate in normal kidney, analyses were restricted to 15 UK laser-capture microdissection (LCM) donors without a history of kidney cancer or overt renal pathology, thereby minimizing confounding from geographic variation and disease status and providing a baseline representative of the general population. Model selection was guided by likelihood-ratio tests comparing nested models to determine the inclusion of fixed and random effects. Final models included age–structure interaction terms as fixed effects, with patient-specific random slopes to account for within-patient correlation. Heteroscedastic residual variance across nephron structures was modelled using structure-specific variance functions, and models were fitted by restricted maximum likelihood.

For single-base substitution signatures contributing to more than 1% of the total mutation burden, structure-specific Samples from the same structure within the same donor were pooled to reduce technical noise. Models included age–structure interactions as fixed effects and random intercepts for country and donors nested within country. Regionally restricted signatures (SBS22a, SBS22b, SBS22c, SBS12) were excluded from rate estimation because they are associated with episodic exposures rather than continuous age-related accumulation.

### Mutational signatures associated with smoking, sex, age

Associations between mutational signature burden and donor characteristics were assessed using linear mixed-effects models within the same general framework: Age was included as a fixed effect in all models. Sex and tobacco exposure were included where biologically relevant (tobacco for SBSB, ID3 and DBS2). Country was modelled as a random intercept to control geographic correlation.

For each covariate of interest, statistical significance was assessed using likelihood ratio tests comparing full and reduced models. *P* values were adjusted across signatures using the Benjamini–Hochberg (BH) procedure. All tests were two-sided.

### Mutational signatures associated with cancer incidence

To assess whether inter-country variation in cancer incidence is associated with mutational signature burden, we estimated country-specific signature burdens at a common reference age (median age=62) and correlated these estimates with age-standardized cancer incidence (ASR).

For mutational signatures reported to be associated with cancer incidence (SBS40a, SBS40b, ID5, ID8), linear mixed-effects models were fitted to individual normal kidney cortex samples, modeling signature burden as a function of age with random intercepts and random age slopes for country. Age was centered at the cohort median (62 years) so that model intercepts corresponded to predicted signature burden at the median age. Sex was included as a fixed-effect covariate for signatures with known sex-specific differences (SBS40b, ID5). Models were fitted using restricted maximum likelihood estimation.

Country-level predicted signature burden at the median age was obtained by combining fixed and country-specific random intercept terms. These country-level burden estimates were then associated with country-level ASR using Spearman rank correlation. Multiple testing across signatures was controlled using the BH procedure.

### Geographic variation of mutational signatures

Geographic effects on signature burden were assessed by modeling country as the primary explanatory factor while adjusting for age and, where appropriate, sex or tobacco exposure. For proximal tubules, linear mixed effect models were used to account for multiple samples obtained from the same individual. From each model, estimated marginal means (EMMs) were computed using emmeans (v2.0.1) R package, providing covariate-adjusted country-level burdens.

Country-specific enrichment or depletion was tested using effect-coded contrasts comparing each country to the grand mean. *P* values were adjusted globally across all country–signature comparisons using the BH procedure.

### Variation of mutational signatures in different nephron structures and cancer

To evaluate variation in signature burden across nephron structures and between normal and cancer tissue, we fitted linear mixed-effects models using restricted maximum likelihood (REML). Signature burden was modelled as a function of age and structure, with additional adjustment for sex or tobacco exposure where relevant. Random intercepts for patients nested within country accounted for repeated sampling.

Structure-specific adjusted burdens were estimated using EMMs. Enrichment or depletion of individual structures relative to the overall mean was assessed using effect-coded contrasts, with BH correction applied across all signature–structure comparisons.

## Data availability

Sequencing data have been deposited in the European Genome-phenome Archive (EGA) under accession numbers EGAD00001015825 (for bulk kidney cortex), EGAD00001015827 (for LCM biopsies), EGAD00001015824 (for blood), EGAD00001015826 (for additional cancer whole-genome sequencing) and EGAD00001015828 (for matched normal). The guidelines for patient consent prevent the derived data files from being dispersed by open access. To ensure the data is used for academic and research purposes, controlled access of the data will be available indefinitely upon request made to the WTSI CGP Data access committee.

## Code availability

All software used for data analysis are publicly available with repositories noted within the respective method sections. The bespoke code used for the analyses is available on GitHub (https://github.com/YichenWang1/normal_kidney).

## Acknowledgement

The authors thank L. O’Neill, K. Roberts, K. Smith, S. Austin-Guest and the staff of Sequencing Operations at the Wellcome Sanger Institute for their contribution; IARC General Services, including the Laboratory Services and Biobank team led by Z. Kozlakidis and the Section of Support to Research overseen by C. Mehta, under IARC regular budget funding; the Leeds Biobanking and Sample Processing Lab, the Leeds Multidisciplinary RTB, the Leeds NIHR BioRTB and V. Janout from Palacky University for provision of samples. M. Clatworthy, N. McGranahan, P. Nicola and E. Dunstone for useful discussions, and all the patients involved in this study and their families. This work was supported by the Wellcome Trust grants 206194 and 220540/Z/20/A. Some of this work was delivered as part of the Mutographs team supported by the Cancer Grand Challenges partnership funded by Cancer Research UK (C98/A24032). The sample collection was also partly supported by Barretos Cancer Hospital, the Public Ministry of Labor of Campinas (Research, Prevention, and Education of Occupational Cancer, 2015 to R.M.R.), and by Hospital de Clínicas de Porto Alegre (180330 to B.S.N.), the Practical Research Project for Innovative Cancer Control from the Japan Agency for Medical Research and Development (AMED) (JP20ck0106547h0001 to T.S.), the 1st and 2nd Faculties of Medicine, Charles University, Prague (CAGEKID to I.H.; Occupation, Environment and Kidney Cancer in Central and Eastern Europe to A.H.), the Ministry of Health of the Czech Republic (MH CZ–DRO (MMCI, 00209805) to L.F. and M.N.), NIHR GOSH Biomedical Research Centre and NIHR Cambridge Biomedical Research Centre (NIHR203312 to S.B.). Y. Wang and M. H. Pham were funded by Wellcome Ph.D. studentships. S.B. receives a personal Fellowship from Wellcome (reference 223135/Z/21/Z). The views expressed are those of the authors and not necessarily those of their funders. For the purpose of open access, the authors have applied a CC-BY public copyright license to any author-accepted manuscript version arising from this submission.

## Author contributions

M.R.S. and Y.W. conceptualized the project. Y.W. led the data analysis with support from F.A., M.H.P., P.J.C., S.M., C.N., and S.F. Y.W. and W.K. led the experimental work with support from Y.H., M.H.P. and C.D. Pathology review was carried out by B.A.-A. F.A., I.M. and A.R.J.L. contributed to method development. T.C. and V.G contributed to data curation. Scientific project management was carried out by L.H., A.F.-I., and S.P. Patient and sample recruitment was led or facilitated by A.F.-I, A.F., T.S., V.J., S.R., D.G.Z., A.F.M., C. Sima., I.H., A.H., N.S.V., R.E.B., S.O., S. Savic, M.P.C., S.d.C.Z., R.M.R., W.J.M., F.V., B.S.N., S.J., A.Z., L.F., M.N., L.P., C. Shire, W.A. and S. Sangkhathat, K.S.-P, S.B. and C. Latimer. M.R.S., P.B., L.B.A., and P.J.C provided supervision. M.R.S. and Y.W. wrote the manuscript, and all authors contributed to reviewing and editing it.

---

**Extended Data Fig.1.**
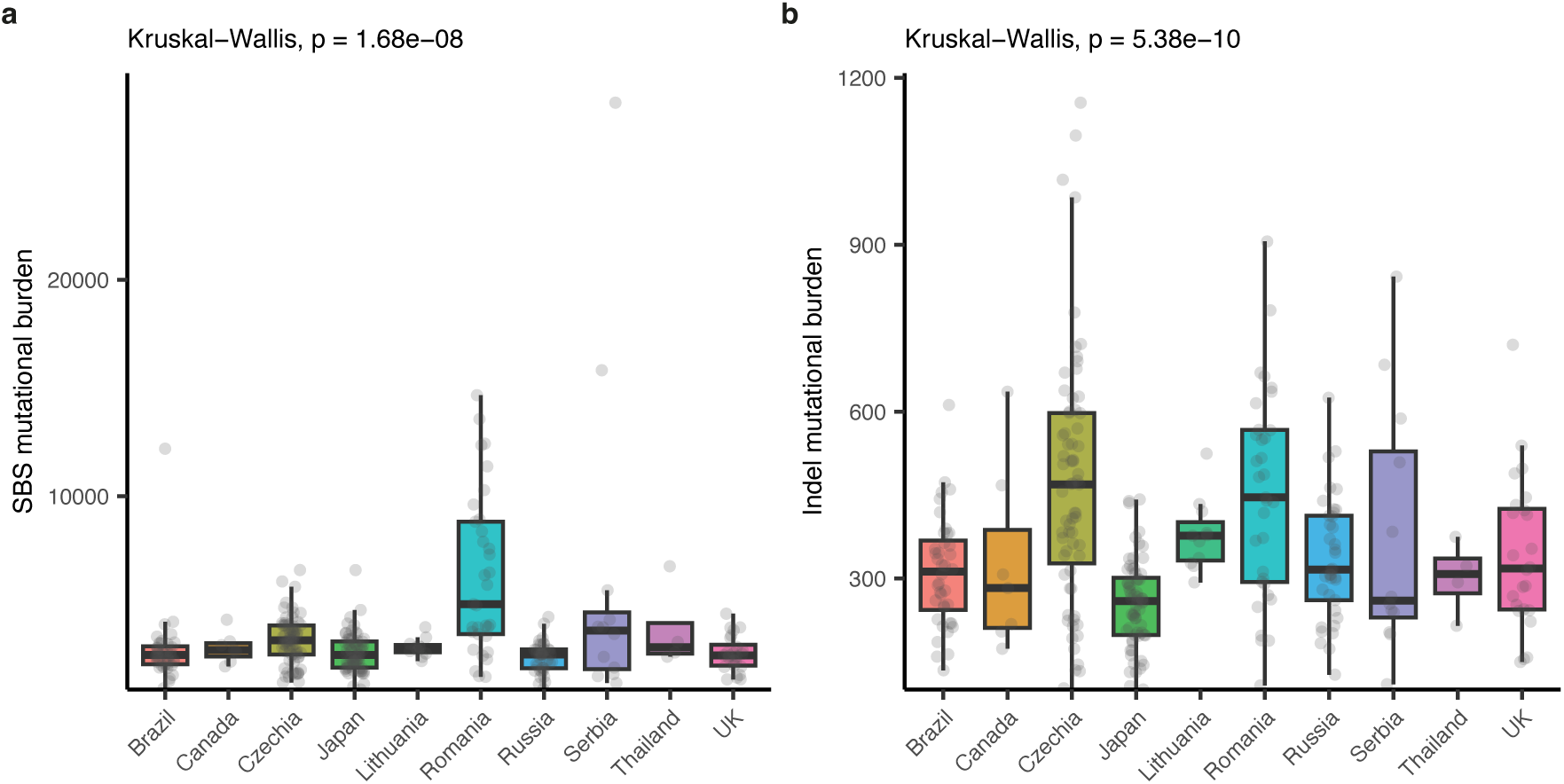
Mutation burdens in normal kidney cortex across countries. Mutation burdens for single base substitutions (SBS) **a,** and small insertions and deletions (indel) **b,** show significant differences between countries using the Kruskal-Wallis (two-sided) test. A total of 299 biologically independent samples are shown: Brazil (*n* = 40), Canada (*n* = 7), Czechia (*n* = 67), Japan (*n* = 64), Lithuania (*n* = 11), Romania (*n* = 33), Russia (*n* =37), Serbia (*n* = 12), Thailand (*n* = 4), UK (*n* = 24).The central line, box and whiskers represent the median, the 25th and 75th percentiles, and 1.5 × interquartile range respectively.

**Extended Data Fig.2.**
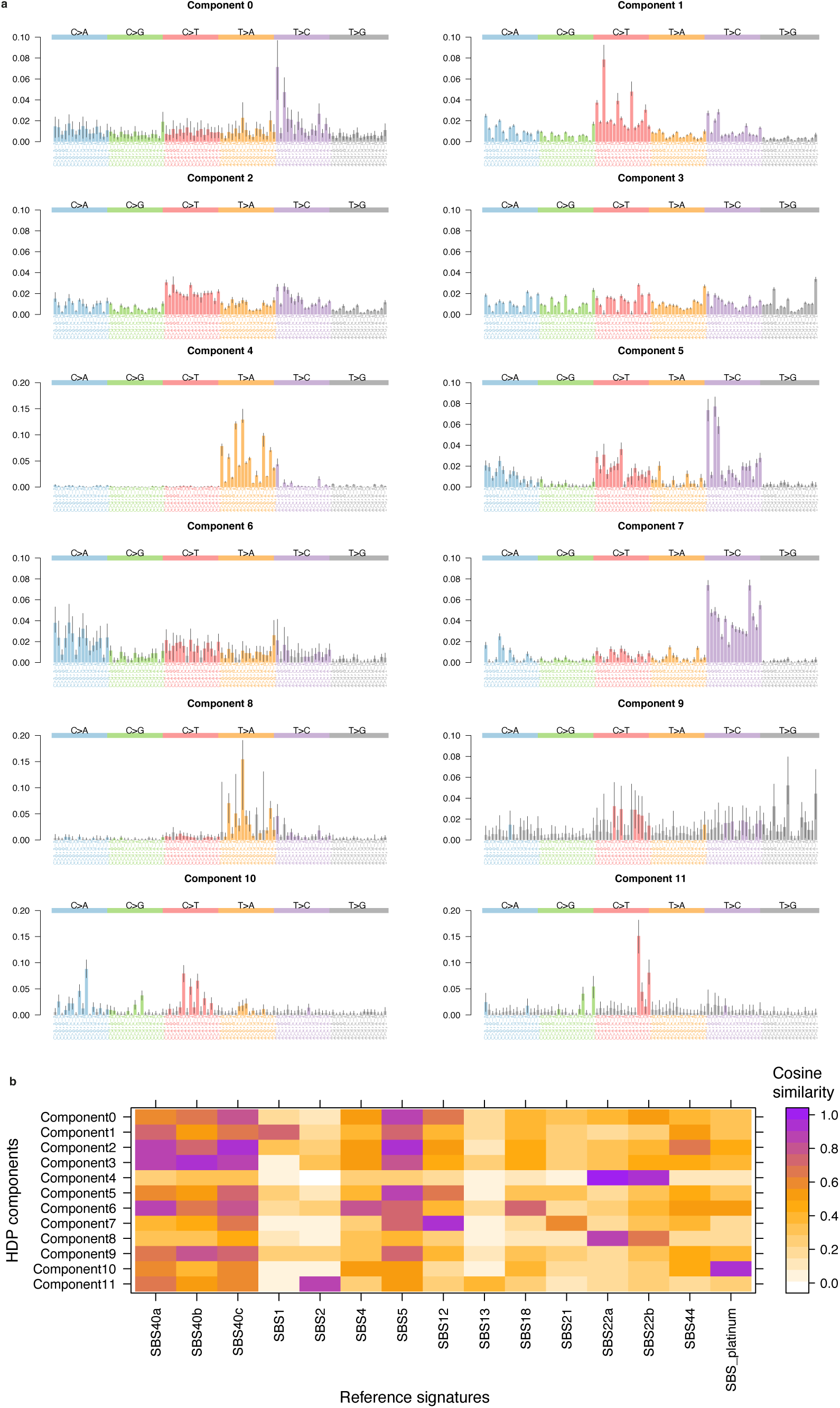
Single-base substitution signatures from *de novo* HDP extraction. **a**, Extracted HDP single-base substitution signatures. X axis represents 96 categories of trinucleotide context and y axis represents the proportion. The black lines indicate 95% confidence intervals. Trinucleotide contexts that are not statistically significant are shown in light grey. **b**, Cosine similarity between HDP signatures and mutational signatures reported in kidney cancer.

**Extended Data Fig.3.**
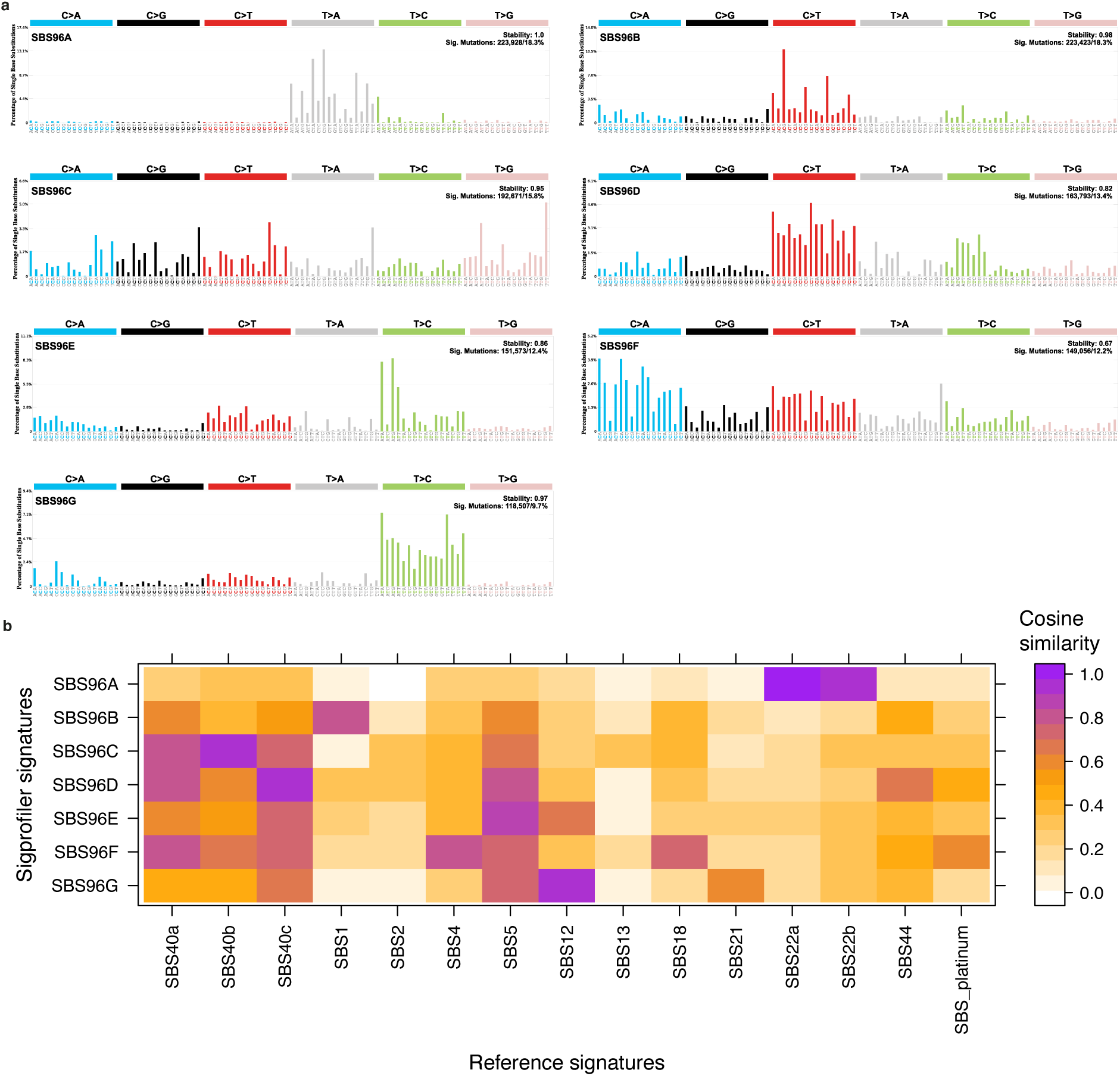
Single-base substitution signature from *de novo* SigProfiler extraction. **a**, Extracted SigProfiler single-base substitution signatures. X axis represents 96 categories of trinucleotide context and y axis represents the proportion. **b**, Cosine similarity between SigProfiler signatures and mutational signatures reported in kidney cancer.

**Extended Data Fig.4.**
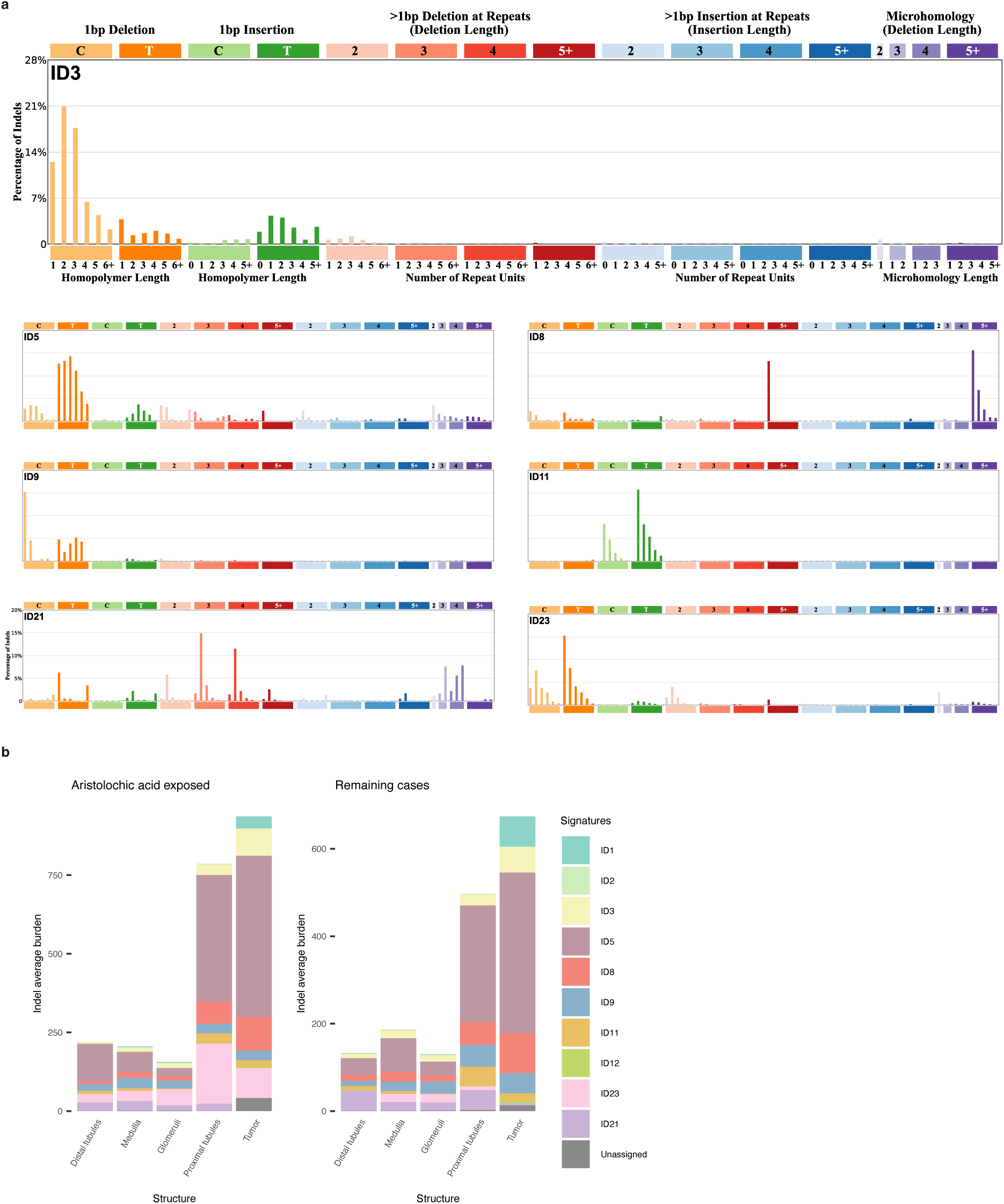
Indel mutational signatures across different kidney structures. **a**, Indel mutational signatures in normal kidneys. **b**, Average attribution for indel signatures across different nephron structures in ccRCC patients and corresponding cancer genomes. Left, aristolochic-acid exposed donors (*n* = 14). Right, all remaining donors (*n* = 54).

**Extended Data Fig.5.**
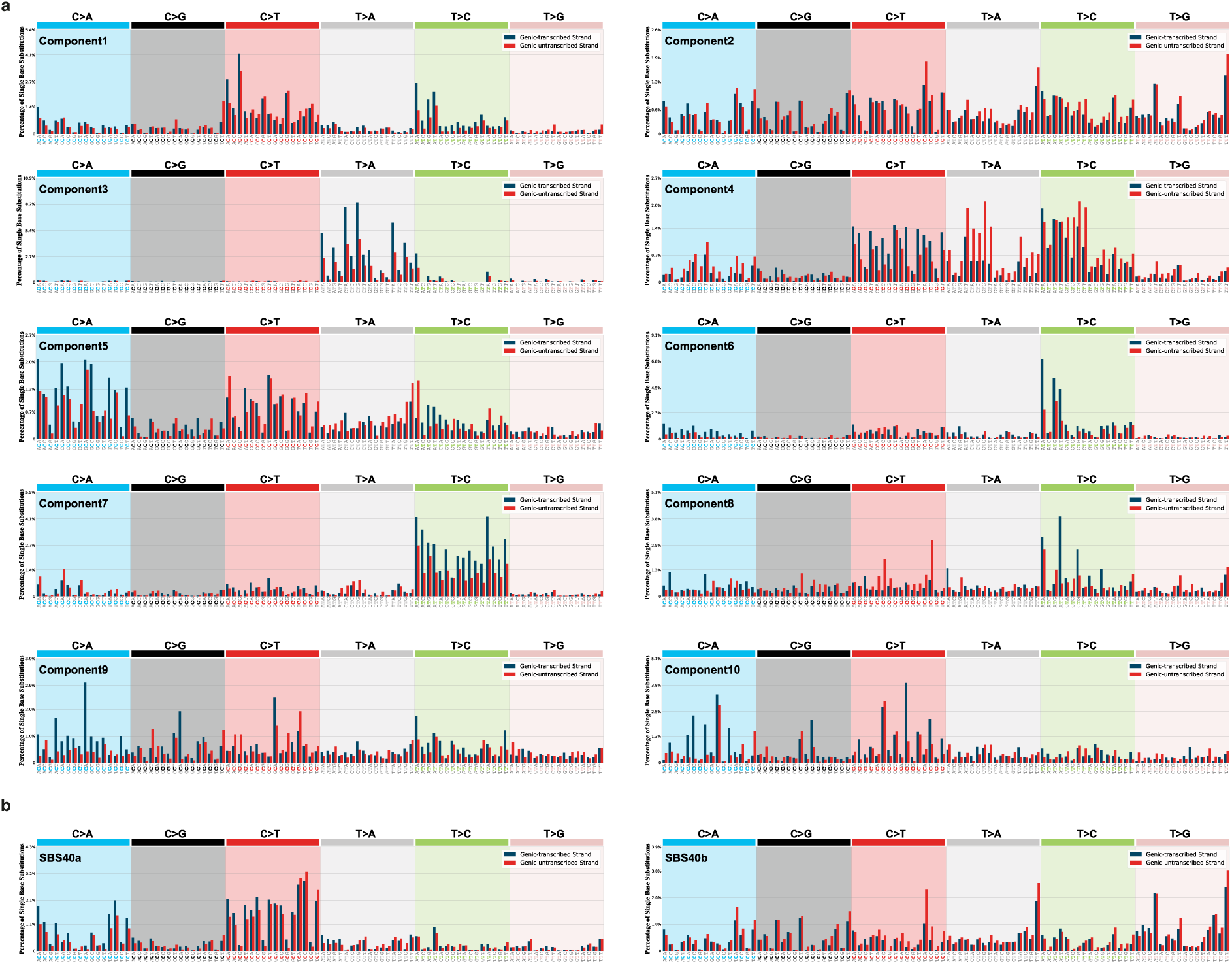
*De novo* extraction of SBS288 signature showing transcriptional strand bias. **a**, Mutational signatures extracted in normal kidney dataset by HDP using 288 categories (96 categories of trinucleotide context in x axis in combination with transcribed strand, non-transcribed strand or intergenic region). **b**, Further splitting of SBS40a and SBS40b when extracted with cancer samples, showing their transcriptional strand preference in different trinucleotide contexts.

**Extended Data Fig.6.**
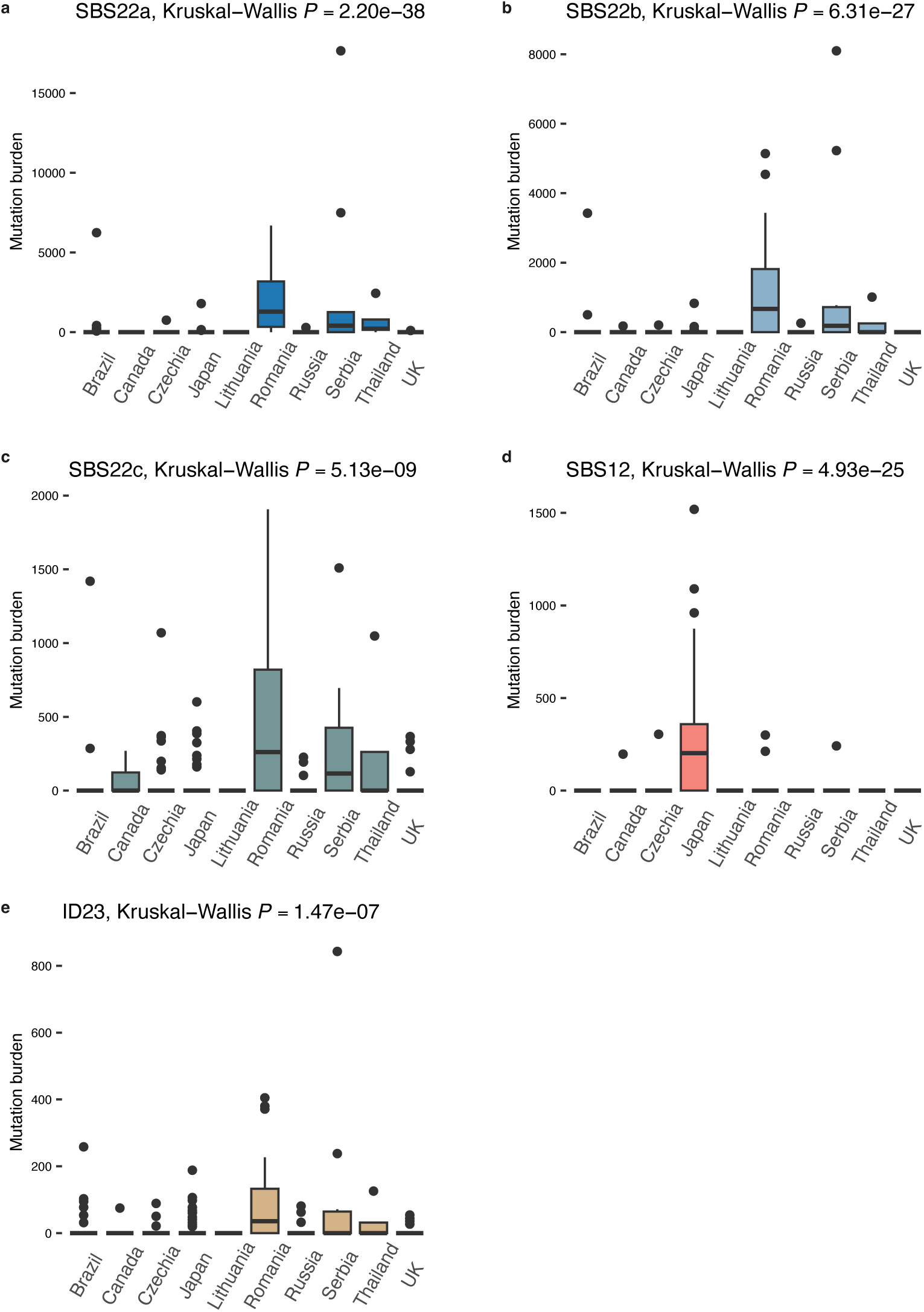
Distribution of region-specific signatures. Mutation burdens for single base substitution signature SBS22a (**a**), SBS22b (**b**), SBS22b (**c**), SBS12 (**d**) and indel signature ID23 (**e**) in different countries. *N* = 299 from Brazil (*n* = 40), Canada (*n* = 7), Czechia (*n* = 67), Japan (*n* = 64), Lithuania (*n* = 11), Romania (*n* = 33), Russia (*n* =37), Serbia (*n* = 12), Thailand (*n* = 4), UK (*n* = 24). The central line, box and whiskers represent the median, the 25th and 75th percentiles, and 1.5 × interquartile range respectively.

**Extended Data Fig.7.**
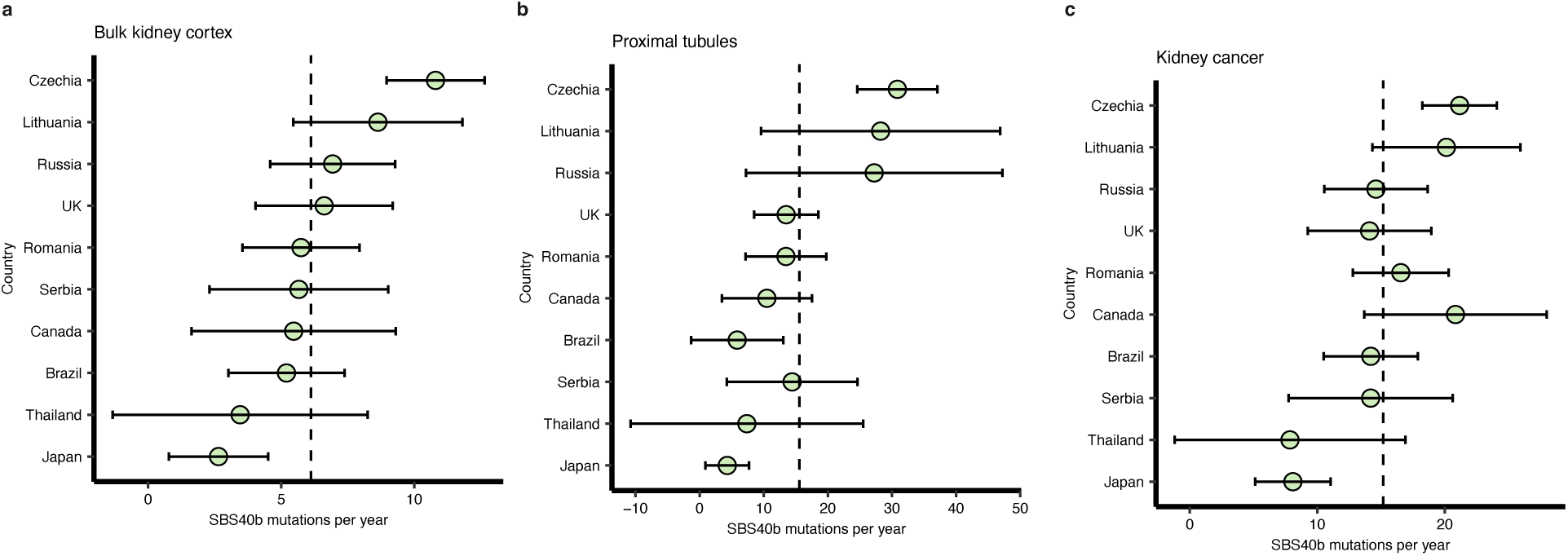
Estimated SBS40b mutation rates across different countries. Mutation rates estimated from bulk kidney cortex (**a**), proximal tubules (**b**) and corresponding cancer genomes(**c**). Central dots represent the estimated fixed effect from linear mixed-effects models, and error bars represent the 95% confidence intervals. Dash lines represent average mutation rate across all countries investigated. *N* = 299 for bulk kidney cortex, 166 for proximal tubules, 291 for cancer.

**Extended Data Table 1.** Previously reported somatic mutation rates in healthy normal human tissues. Mutation rates are estimated per diploid genome per year.

| Tissue | Rate (SNVs/yr) | Rate (Indels/yr) |
| --- | --- | --- |
| Skin (sun-exposed)<sup>9</sup> | Variable, can reach >200 | Not reported |
| Bronchial epithelium (current smoker)<sup>10</sup> | Variable, can reach >100 | Variable, can reach >2 |
| Colorectal epithelium<sup>34</sup> | ~44 | ~1–2 |
| Small intestinal epithelium<sup>26</sup> | ~42–51 | ~2–4 |
| Esophagus<sup>38,39</sup> | ~42 | Not reported |
| Hippocampal dentate gyrus neurons<sup>40</sup> | ~40 | Not reported |
| Bladder epithelium<sup>27</sup> | ~10–30 | Not reported |
| Normal bronchial epithelium (non-smoker)<sup>10</sup> | ~22 | ~1 |
| Gastric epithelium<sup>37</sup> | ~28 | ~2 |
| Endometrial epithelium<sup>41</sup> | ~29 | ~1–2 |
| Liver hepatocytes<sup>36</sup> | ~33 | ~1–2 |
| Lymphocytes<sup>42</sup> | ~15–25 | ~0.8–1 |
| Hematopoietic stem cells<sup>43</sup> | ~17 | ~0.7 |
| Frontal cortex neurons<sup>14,40</sup> | ~17–23 | ~2.5 |
| Smooth muscle<sup>14</sup> | ~21 | ~1.3 |
| Prostate<sup>22</sup> | ~19 | Not reported |
| Pancreatic acini<sup>22</sup> | ~15 | Not reported |
| Bile duct<sup>22</sup> | ~9 | Not reported |
| Seminiferous tubules<sup>22</sup> | ~2.7 | Not reported |

